# Generation of a biosafety mouse model infection for SARS-CoV-2 replicon delivery particles

**DOI:** 10.1101/2024.11.10.622838

**Authors:** Yingjian Li, Xiaoya Huang, Jikai Deng, Xue Tan, Qianyun Liu, Li Zhou, Yu Chen

## Abstract

Animal models are essential for understanding the pathogenesis of SARS-CoV-2 and for developing therapeutic strategies. Replicon delivery particles (RDPs) were a component of trans-complementary systems for SARS-CoV-2, which is a safe and convenient tool in researching SARS-CoV-2 in animal biosafety level-II laboratory (ABSL-2). Here, we constructed a mouse model that conditional expressing SARS-CoV-2 N on the background of the K18-hACE2 KI mice. The SARS-CoV-2 N with flanked loxP-stop-loxP sequence was under the CAG promoter, and this cassette was knocked into the Tiger locus of mouse by CRISPR-Cas9 (K18-hACE2-N KI). By mating K18-hACE2-N KI mice with Cre tool mice, the offspring can express SARS-CoV-2 N (Cre-N-hACE2 KI) systemically and in a tissue-specific manner. Cre-N-hACE2 KI exhibited susceptibility to the SARS-CoV-2 ΔN-GFP/HBiT infection. The viral loads in lung exhibited a mountain-like trend, peaking at 4 days post-infection, and lung injuries can be observed. Overall, we demonstrated a mouse model infection for SARS-CoV-2 ΔN-GFP/HBiT to understand SARS-CoV-2 pathogenesis.

## Introduction

The pandemic caused by severe acute respiratory syndrome coronavirus 2 (SARS-CoV-2) has taken millions of lives and is still a threat to global health ^1,2^. Coronaviruses are also responsible for another two epidemics over the past two decades, severe acute respiratory syndrome coronavirus (SARS-CoV-2) in 2003 and Middle East respiratory coronavirus (MERS-CoV) in 2012 ^3,4^. As the SARS-CoV-2 variants emerging, SARS-CoV-2 infection usually results in mild symptoms. However, for immune-compromised people, symptoms can be developed to severe disease with acute lung injuries and acute respiratory distress syndrome (ARDS), and even to death ^2,5^. It is reported that COVID-19 death tends to be related with the “cytokines storm” characterized with the increased levels of some cytokines ^6^.

Small animal models are indispensable for SARS-CoV-2 research. SARS-CoV-2 takes human angiotensin-converting enzyme 2 (hACE2) as receptor for viral entry, and natural mouse is not susceptible to SARS-CoV-2 due to the ineffective binding between SARS-CoV-2 spike protein and murine ACE2 ortholog (mACE2) ^3,7^. Several transgenic mouse lines expressing hACE2 models have been developed so far. These model generation strategies encompass the expression of hACE2 under the control of exogenous or mACE2 promoters, as well as the transduction of hACE2 utilizing adenoviral vectors ^8-12^. Additionally, some groups have employed mouse-adapted SARS-CoV-2 strains that can infect wild-type mice, circumventing the need for hACE2 expression^9,13,14^. Each mouse model has its own characteristics and advantages, with symptoms ranging from mild to severe, and even lethal. The mouse model can mimic the lung injuries that observed in the COVID-19 patients, providing a valuable tool for studying the pathophysiological mechanism of the disease and assessing the potential antiviral drugs and vaccines ^15,16^.

Concerning the biosafety, replicon and trans-complementary system offer convenient substitutions for some highly pathogenic viruses, such as Ebola and coronaviruses ^17-20^, thereby lowering the threshold of research on these viruses. The trans-complementation system for SARS-CoV-2 consists of a genomic viral RNA containing a deletion of structure gene and a producer cell line expressing the deleted gene^21^. Trans-complementation of the two components generates a virion that can mimic the life cycle of virus in the permissible cells. The trans-complementary system of SARS-CoV-2 has proven valuable for facilitating the study of the SARS-CoV-2 and high-through evaluation of the antiviral drugs. Several groups have also established the corresponding mouse model for trans-complementation derived virions infection^22^.

In this study, we established a SARS-CoV-2 ΔN-GFP/HBiT infection mouse model on the K18-hACE2 KI mouse background, which expresses the SARS-CoV-2 N. Combined with tissue-specific Cre tool mice, two types of mice with systemic and lung-specific expression of SARS-CoV-2 N protein were generated. We then evaluated the susceptibility, viral replication, pathological symptoms.

## Results

### The construction of SARS-CoV-2 N conditional knock-in mouse on the background of K18-hACE2 KI mouse

Based on the principle of constructed SARS-CoV-2 trans-complementary system, we considered to generate a mouse expressing hACE2 and SARS-CoV-2 N protein. There have been several reported mice expressing hACE2 for authentic SARS-CoV-2 infection, and we are more similar and preferred to the K18-hACE2 KI mouse among those mice. So, we designed and constructed the SARS-CoV-2 N conditional knock-in mouse on the background of this mouse strain. CAG-loxP-stop-loxP-Kozak-SARS-CoV-2 N-WPRE-polyA cassette was inserted into the Tigre locus on mouse chromosome 9 using CRISPR-Cas9 (Fig 1A). The resulting F0 mice were genotyped (Fig1B), and the target mice were then bred with the K18 hACE2 KI mice to obtain the homozygous K18-hACE2-N KI mice. The Cre-N-hACE2 KI mice were obtained by mating K18-hACE2-N KI mice with Cre tool mice (Sftpc-IRES-iCre or Rosa26-SA-CreERT2 mice). All primers used for mice genotyping were listed in the Table S1.

**Figure 1.**
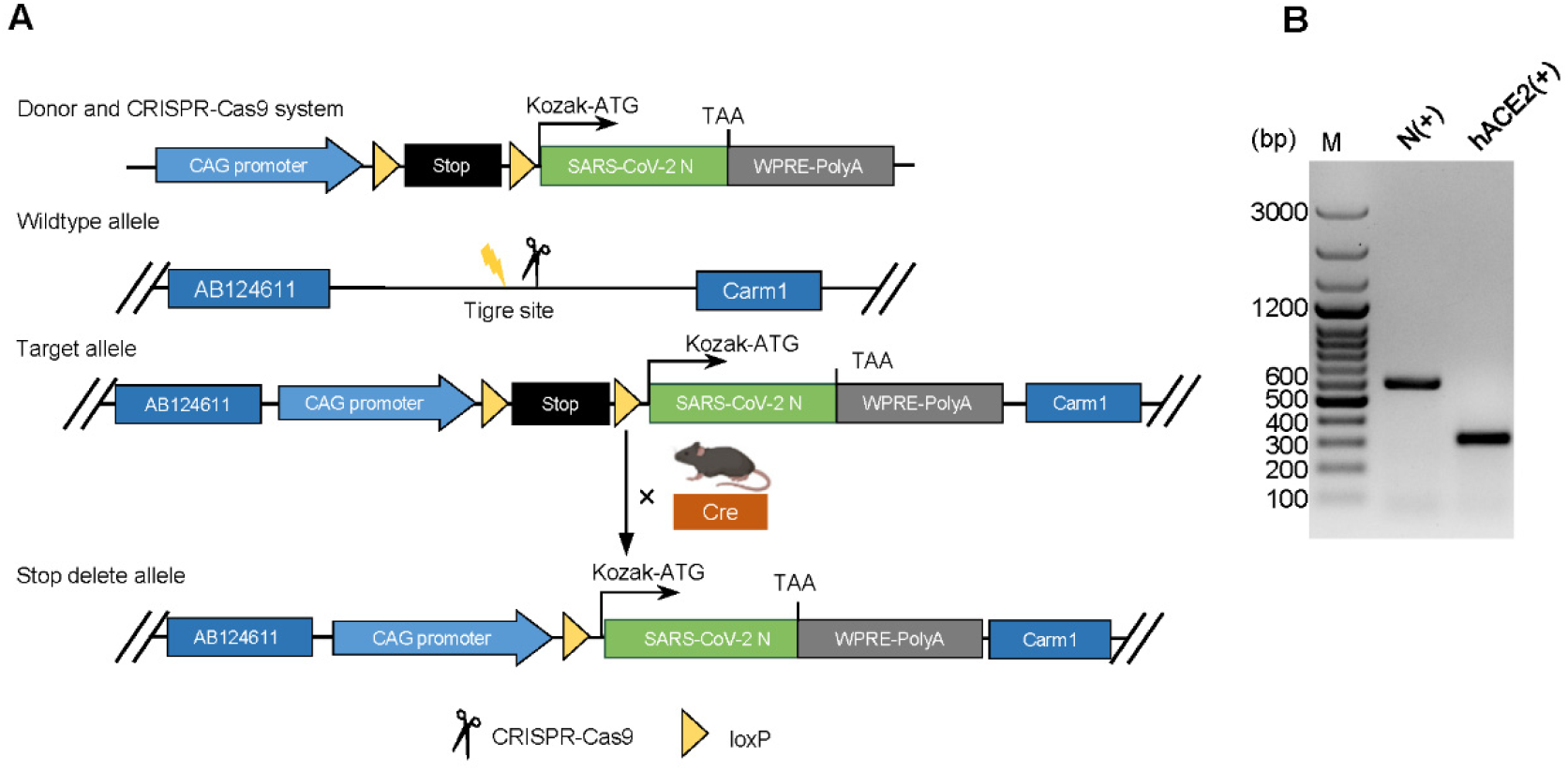
The construction of SARS-CoV-2-N conditional knock in mouse. (A) Schematic diagrams illustrating the knock in strategy. The CAG-LSL-SARS-CoV-2 N-WPRE-PolyA sequence were inserted into Tigre locus on chromosomes 9 on the background of K18-hACE2 KI mouse. When Cre recombinase expressed, the stop sequence between the loxP will be deleted. (B) The genotyping of Tigre-N mouse. The identified fragments of N gene and hACE2 were indicated. M: 100 bp plus DNA ladder.

### SA-N-hACE2 mice are susceptible to SARS-CoV-2 ΔN-GFP/HiBiT

SA-N-hACE2 KI mice were received with 75 mg/kg tamoxifen (TAM) every 24 h for 5 days, and fed for another 7 days for SARS-CoV-2 N protein expression. It showed that SARS-CoV-2 N were detected in multiple tissues of SA-N-hACE2 KI after TAM induction (Fig 2A). The immunofluorescence (IF) analysis further confirmed the expression of N protein in the lungs and demonstrated the expression of hACE2 protein (Fig 2B). For mice challenge experiment (Fig 2C), we firstly determined the viral distribution among tissues. TAM-induced SA-N-hACE2 mice were intranasally inoculated (i.n) with 1×10^6^ TCID_50_ SARS-CoV-2 ΔN-GFP/HiBiT and tissues were collected at 7 days post-infection (dpi). Viral loads can be detected in the brains and lungs (Fig 2D). Then, to further investigate the characteristics of the SA-N-hACE2 mice, mice were i.n with 5×10^4^ TCID_50_ (low does) and 1×10^6^ TCID_50_ (high does) SARS-CoV-2 ΔN-GFP/HiBiT, respectively. The mice were weighted daily and tissue samples were collected at indicated time. Both high-does and low-does group mice lost slight weight (Fig 2E). It demonstrated that viral loads in lung rise within first 4 dpi and fall between 4 to 7 dpi, reaching peak at 4 dpi (Fig 2F). And the high-does infected mice exhibit higher viral loads in the lungs compared to those in the low-does infected mice.

**Figure 2.**
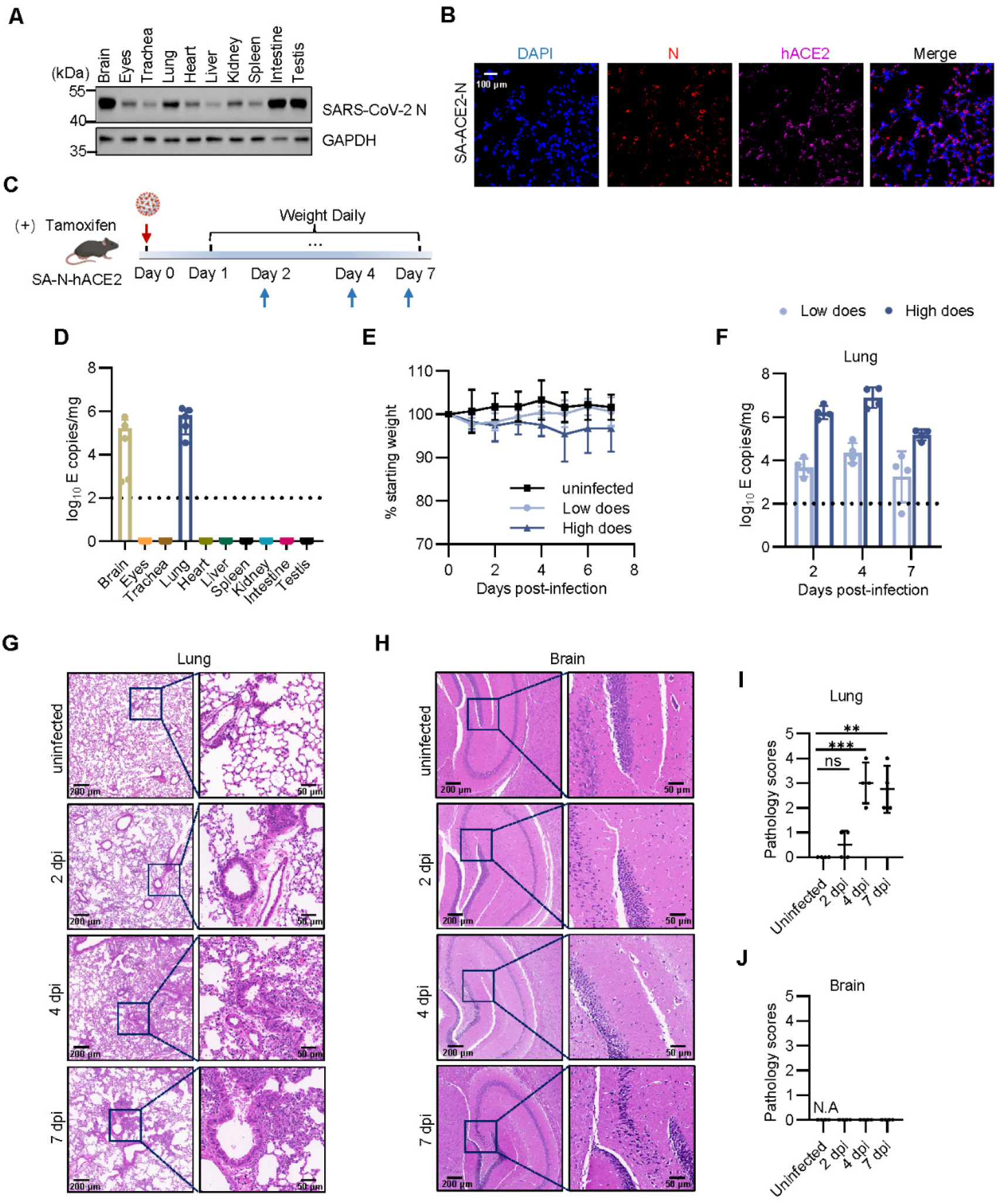
SA-N-hACE2 mice are susceptible to SARS-CoV-2 ΔN-GFP/HiBiT. (A) The expression for SARS-CoV-2 N protein among the tissues of tamoxifen-induced in SA-N-hACE2 mice. (B) The IFA analysis of SARS-CoV-2 N and hACE2 protein in the lung of SA-N-hACE2 mice. (C) The scheme illustrating the intranasal infection. The tissue samples were collected at indicated days post-infection (dpi). (D) The viral loads among the tissues collected at 7 dpi were quantified using RT-qPCR. (E) The mice weight change. (F) The viral load in lungs collected at 2, ACE2 mice were infected with 1× 10^6^ / 4 × 10^4^ TCID_50_ SARS-CoV-2 ΔN-GFP-HiBiT; n = 4 mice per group. (G) Pathological changes in lungs and brains (H) of 1×10^6^ TCID_50_ infected SA-hACE2-N mice at 0,2,4 and 7 dpi. (I) The pathology scores of lungs and (J) brains.

We performed the histopathological analysis on the infected lung and brain tissues from high-does group. It indicated that the lung exhibited a gradual progression of pneumonia, from mild to severe (Fig 2G and 2I). At 2 dpi, there were slight damage in the lungs of the mice. At 4 and 7 dpi, the lungs exhibited severe tissue damage characterized by an increased infiltration of immune cells and the thickening of the alveolar walls. Additionally, no pathological changes were observed in the brains of the high-does infected mice when compared to those of uninfected mice (Fig 2H and 2J).

### Sftpc-N-hACE2 mice exhibit lung-specific susceptibility to SARS-CoV-2 ΔN-GFP/HiBiT

We initially detected the expression of the N protein in Sftpc-N-hACE2 mice. The result showed that the N protein is expressed exclusively in the lung, which is consistent with the expectation (Fig 3A). The immunofluorescence analysis also demonstrated the N and hACE2 protein expression in the lung (Fig 3B). When Sftpc-N-hACE2 mice were i.n infected with 1×10^6^ TCID_50_ SARS-CoV-2 ΔN-GFP/HiBiT, the viral loads were detected solely in the lungs at 7 dpi (Fig 3D), which suggests a correlation between viral loads and the expression of N protein.

**Figure 3.**
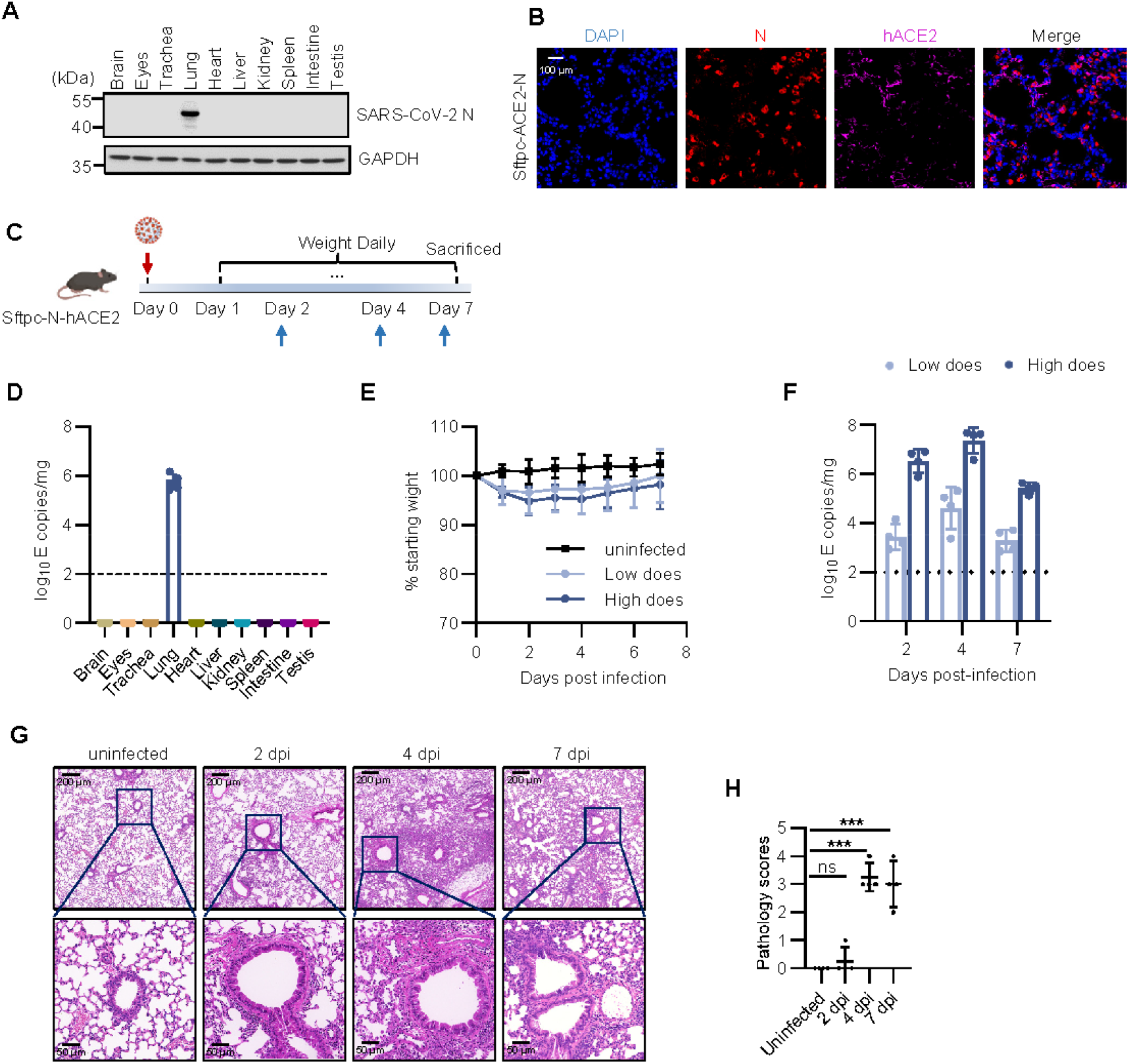
Sftpc-N-hACE2 mice exhibit lung-specific susceptibility to SARS-CoV-2 ΔN-GFP-HiBiT. (A) The expression for SARS-CoV-2 N protein among the tissues of tamoxifen-induced in Sftpc-N-hACE2 mice. (B) The IFA analysis of SARS-CoV-2 N and hACE2 protein in the lung of SA-N-hACE2 mice. (C) The scheme illustrating the intranasal infection. The tissue samples were collected at indicated days post-infection (dpi). (D) The viral load among the tissues collected at 7 dpi were quantified using RT-qPCR. (E) The mice weight change. (F) The viral load in lungs collected at 2, 4 and 7 dpi. The Sftpc-N-hACE2 mice were infected with 1×10^6^ / 4 × 10^4^ TCID_50_ SARS-CoV-2 ΔN-GFP-HiBiT; n = 4 mice per group. (G) Pathological changes in lungs of Sftpc-N-hACE2 mice at 2, 4 and 7 dpi. (H) Pathology scores of lungs.

As the SA-N-hACE2 mice, same experiments were also conducted to elucidate the characteristics of the Sftpc-N-hACE2 mice post infection (Fig 3C). Both low-does and high-does infected mice lost slight weight at first 3-4 dpi (Fig 3E). The viral loads in lung of low-does and high-does infected mice increased within first 4 dpi and then declined (Fig 3F), which exhibit similar trend that observed in the lungs of SA-N-hACE2 mice. Meanwhile, it indicated that mice infected with high does virus exhibit higher viral loads in lung than those in low-does group. High-does group exhibited only mild pathological changes in lung at 2 dpi, such as limited infiltration of inflammatory cells and slight thickening of the bronchial walls. As the infection progresses, the lung damage become much severe at 4 and 7 dpi, characterized by the collapsed epithelial cells, thickened alveolar walls, alveolar damage and infiltration of inflammatory cells around the pulmonary blood vessels (Fig G-H).

## Discussion

Mouse models are most frequently used in understanding the pathogenesis of SARS-CoV-2 and in exploring therapeutic strategies for COVID-19. Poor binding capacity of mouse ACE2 to viral S protein has prompted researchers to implemented several strategies to enhance the mice’s susceptibility to SARS-CoV-2. Generation of transgenic mice expressing hACE2 was one of primary approaches. These transgenic mouse models have been generated by expressing hACE2 under the control of various promoters, such as K18 promoter and the endogenous mouse ACE2 promoter. Meanwhile, development of mouse-adapted of SARS-CoV-2 is another approach that facilitates the research in COVID-19. The major threshold of those mouse models is the requirement in laboratory settings, which is crucial for biosafety. Not only SARS-CoV-2, but also the trans-complementary systems for other highly pathogenic viruses have been increasingly applied in BSL-2. Only a few mouse infection models compatible with the produced viral-like particles from trans-complementation system have been reported.

As we reported previously, we produced replicon delivery particles, SARS-CoV-2 ΔN-GFP/HiBiT, from trans-complementary systems and preliminary results indicate that the replicon delivery particles can replicate in the mouse that expressing both N and hACE2 simultaneously. Here, we introduced a transgenic mouse model that is susceptible to SARS-CoV-2 ΔN-GFP/HiBiT. By selecting different Cre tool mice, SA-N-hACE2 mouse that expresses SARS-CoV-2 N systemically and Sftpc-N-hACE2 mouse that specifically expresses SARS-CoV-2 N in the lung. It showed that those mice are susceptible to SARS-CoV-2 ΔN-GFP/HiBiT. The viral loads were mainly detected in the brain and lung tissues of SA-N-hACE2 mice. By restricting SARS-CoV-2 N to lung-specific expression, viral loads were solely detected in lung tissues of Sftpc-N-hACE2 mice. It is consistent with expectation that SARS-CoV-2 N is required for the viral replication and transcription and the principle of trans-complementary system. The viral load in lung tissues of both SA-N-hACE2 and Sftpc-N-hACE2 mice presented an increasing trend during the first 4 dpi and then drop at 7 dpi. It indicated that the viral replication characteristics in the lung tissues of both mice are similar. High does RDPs infection can induce progressive pneumonia, which featured by epithelial cells, alveolar injury and perivascular inflammatory cell infiltration. Although viral loads can be detected in the brain tissues of SA-N-hACE2 mice, there was no significant damage observed, a phenomenon also seen in several SARS-CoV-2 infected hACE2 transgenic mice models. Meanwhile, we found that some infected SA-N-hACE2 mice exhibited signs of lethargy and a hunched back, which could be related with the brain infection.

In summary, our research established a mouse model that is susceptible to intranasal infection of SARS-CoV-2 ΔN-GFP/HiBiT. Our results demonstrated the pathogenicity of SARS-CoV-2 ΔN-GFP/HiBiT in SA-N-hACE2 and Sftpc-N-hACE2 mice with clinical symptoms, such as acute lung injury characterized by inflammatory cell infiltration and thickened alveolar walls, recapitulating symptoms and pathology in patients with COVID-19. This mice model serves as a valuable tool for deciphering the infectivity and pathogenesis of SARS-CoV-2, offering insights while requiring a more accessible laboratory setting.

## Materials and methods

### Viruses, cells and viral propagation

The SARS-CoV-2 ΔN-GFP/HBiT, replicon delivery particles (RDPs), was constructed as described as previously ^21^. About 20 μg full-length mRNA and 10 μg N mRNA was electroporated into Caco-2-N cells. The recused RDPs were propagated in the Caco-2-N cells and the supernatant was collected at appropriate time. The collected supernatant was then concentrated by ultracentrifugation. The SARS-CoV-2 wild-type strain (IVCAS 6.7512) was provided by the National Virus Resource, Wuhan Institute of Virology, Chinese Academy of Science. The SARS-CoV-2 was produced in the Vero-E6 cells. The RDPs and virus stock were titrated in the corresponding cells.

### Mice

Heterozygous C57BL6/JGpt-H11^em1Cin(K18-hACE2)^Tigre^em1Cin(CAG-LSL-SARS-CoV-2-N-Wpre-PolyA)/^Gpt mice (K18-hACE2-N KI mice, #T058284) were constructed on the background of C57BL/6JGpt-H11^em1Cin(K18-ACE2)/^Gpt mice (K18-hACE2 KI mice, #T037657), by inserting the CAG-LSL-SARS-CoV-2-N-Wpre-PolyA fragment into the Tigre site of mouse through the CRISPR-Cas9 technique. The resulting F0 mice were identified by sequencing and PCR. Those processes aboved were conducted by GemPharmatech (Nanjing, China). The C57BL/6JGpt-Sftpc^em1Cin(IRES-iCre)^/Gpt mice (Sftpc-IRES-iCre mice, #T004715) and C57BL/6JGpt-Rosa26^em1Cin(SA-CreERT2)^/Gpt mice (Rosa26-SA-CreERT2 mice, #T050182) were purchased from GemPharmatech. The K18-hACE2-N mice were bred to the Cre tool mice, and the offspring was genotyped by PCR using the primers indicated in Table S1.

### Western Blotting

Tissues were collected and then homogenized. The homogenates were lysed with RIPA buffer containing protease inhibitor cocktail. Lysates were mixed with loading buffer and boiled for 10 min. After centrifugation, the lysates were electroporated in polyacrylamide gel and transferred to PVDF membrane. The membrane was blocked in 10% skim milk at room temperature for 1 h. The membrane was incubated with primary antibody anti-SARS-CoV-2 N or anti-GAPDH for 2 h and then washed in PBS containing 0.1% Tween 20. The membrane was exposed to secondary antibody for 1 h. Followed by 3 times wash, the membrane was visualized using ChemiDoc Imaging system (Bio-Rad).

### Mouse challenge

Eight-ten-week-old SA-N-hACE2 and Sftpc-N-hACE2 mice were anesthetized with isoflurane and intranasally (i.n.) inoculated with 50 μl DMEM containing the SARS-CoV-2 ΔN-GFP/HBiT (1 × 10^6^ TCID_50_ or 5 × 10^4^ TCID_50_). Mice i.n. inoculated with equal volume of DMEM were considered as a mock control. All the mice were monitored and weighted daily. Mice were euthanized at the indicated time to collect the tissues. The tissues were processed and then stored at -80°C_∘_

### RNA isolation and RT-qPCR

Tissues were weighted and homogenized in 1 ml PBS. Tissue homogenates were clarified by centrifugation at 5000 rpm for 5 min. Total RNA was extracted using Trizol LS Reagent (Invitrogen) according to the manufacturer’s instruction. RNA was reverse transcribed using PrimeScript 1st Strand cDNA synthesis Kit (TAKARA). The viral loads were determined by measuring the SARS-CoV-2 E gene copies using a TaqMan multiplex qPCR kit (Yeasen). The standard curves were generated with a plasmid harbouring SARS-CoV-2 E gene.

### Histopathology analyses

Tissue samples were fixed with 4% paraformaldehyde for at least 48 hours, embedded in paraffin, and cut into sections. The fixed tissue samples were used for H&E staining and indirect immunofluorescence assays (IFAs). Antibody used for IFAs are as follows: anti-hACE2 antibody and anti-SARS-CoV-2 N antibody (catalog: 10108-RP01 and 40143-MM05, SinoBiological). The image information was collected using a Pannoramic MIDI system (3DHISTECH).

### Statistical analyses

The results are shown as the mean ± SD or mean ± SEM. Student’s t-test or one-way/two-way ANOVA with multiple comparison test were used to measure the statistical difference between samples using GraphPad Prism 8. statistically significant differences are shown as follows: *, p < 0.05; **, p < 0.01; ***, p < 0.001; ****, p < 0.0001.

For pathology score evaluation, lung damages such as alveolar wall thickness, hemorrhage and immune cell infiltration were observed by H&E staining under the microscope at a magnification of 10 × filed. Five random fields of each section were scored according to the following criteria: score 0, no damage; score 1, mild damage and the area of damage was less than 25%; score 2, moderate damage and the area of damage was 25-50%; score 3, severe damage and the area of damage was 50-75%; score 4, the area of damage was over 75%. The average score of five fields was the individual H&E score of the lung injury ^23^.

## Acknowledgements

We thank the GemPhamatech Co., Ltd (Nanjing, China) for their great support. This study was supported by grants from the National Key R&D Program of China (2021YFF0702004, and 2021YFA1300801).

## Author Contribution

Yingjian Li: conceptualization, methodology, software, date curation and writing-original draft preparation. Xiaoya Huang: methodology, software and date curation. Jikai Deng: methodology and software. Qianyun Liu: methodology. Li Zhou: methodology, investigation, writing reviewing and editing, supervision and funding acquisition. Yu Chen: investigation, supervision, writing-reviewing and editing, and funding acquisition.

## Figure Legend

**Table S1.**
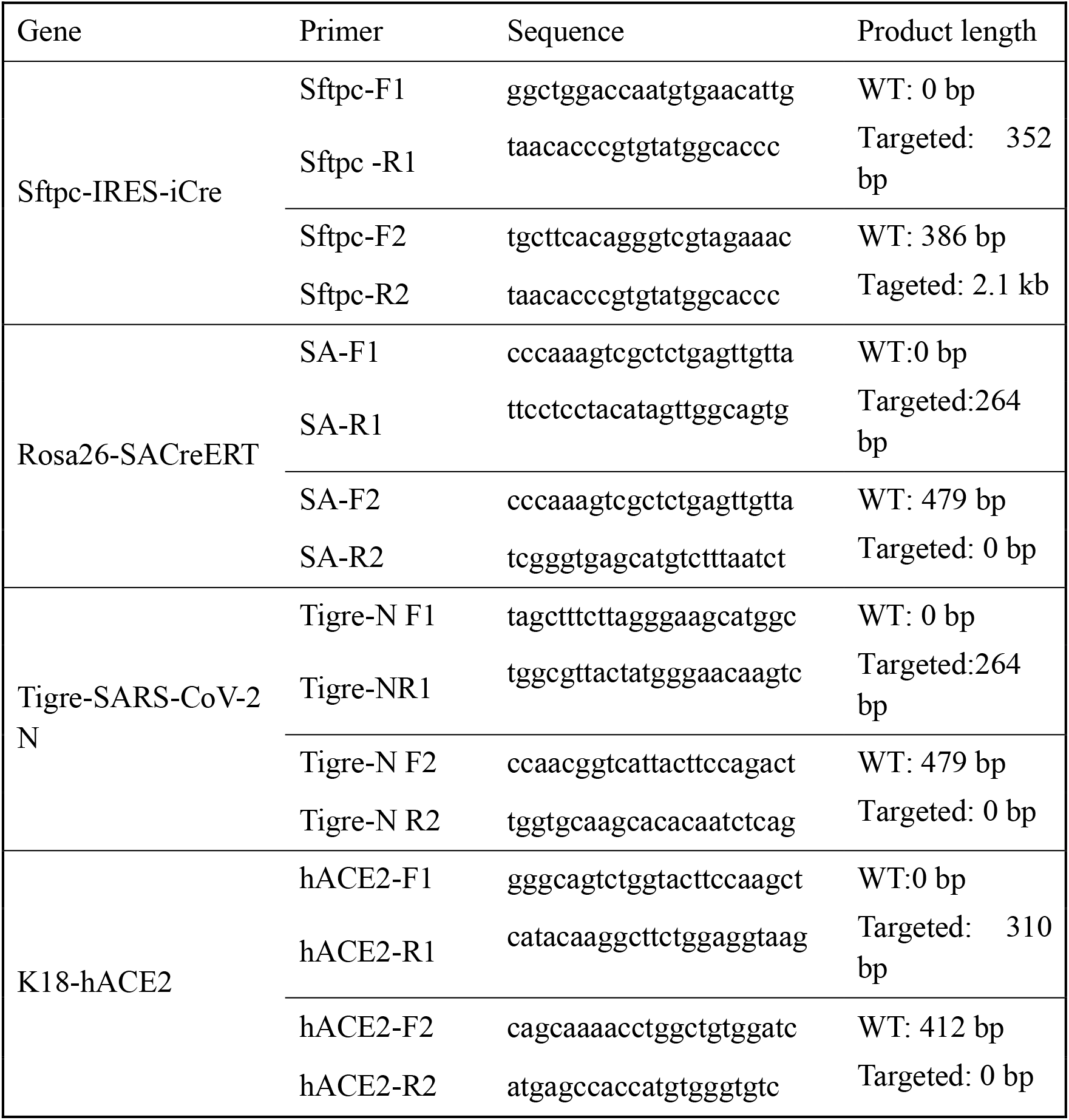
Primers used for mice genotyping.

